# rtPA Directly Protects Neurons After Intracerebral Hemorrhage through PI3K/AKT/mTOR Pathway

**DOI:** 10.1101/2023.02.13.528249

**Authors:** Jie Jing, Shiling Chen, Xuan Wu, Jingfei Yang, Xia Liu, Jiahui Wang, Jingyi Wang, Yunjie Li, Ping Zhang, Zhouping Tang

## Abstract

Intracerebral hemorrhage (ICH) is an acute cerebrovascular disease with high disability and mortality rates. Recombinant tissue plasminogen activator (rtPA) is commonly applied for hematoma evacuation in minimally invasive surgery (MIS) after ICH. However, rtPA may contact directly with brain tissue during MIS procedure, which makes it necessary to discuss the safety of rtPA. We found that, in the in vivo ICH model induced by VII-type collagenase, rtPA treatment improved the neurological function of ICH mice, alleviated the pathological damage and decreased the apoptosis and autophagy level of the peri-hematoma tissue. In the in-vitro model of ICH induced by hemin, the administration of rtPA down-regulated neuronal apoptosis, autophagy, and endoplasmic reticulum stress of neurons. Transcriptome sequencing analysis showed that rtPA treatment upregulated the PI3K/AKT/mTOR pathway in neurons, and PI3K inhibitor (LY294002) can reverse the protective effects of rtPA in inhibiting excessive apoptosis, autophagy and ER-stress. Epidermal growth factor receptor inhibitor (AG-1487) reversed the effect of rtPA on PI3K/AKT/mTOR pathway, which might indicate that the EGF domain played an important role in the activation of PI3K/AKT/mTOR pathway.

## Introduction

Acute spontaneous intracerebral hemorrhage (ICH) is a serious life-threatening acute cerebrovascular disease that accounts for approximately 10%-20% of all strokes (Broderick, Brott et al., 1993, van Asch, Luitse et al., 2010), with a high one-year mortality of 54% and only 12%-39% of survivors achieving long-term functional independence (An, Kim et al., 2017). Irreversible neural damage due to the direct mass effect of the hematoma and the secondary damage induced by hematogenous toxic components may explain the limited effect of present treatment of ICH (Keep, Hua et al., 2012, Zhu, Wang et al., 2019). The early removal of hematoma is the key to the treatment of ICH (Zhou, Wang et al., 2014). Minimally invasive surgery(MIS) plus recombinant tissue plasminogen activator (rtPA) for clot lysis has become a commonly used means of hematoma evacuation after ICH (Hou, Lu et al., 2021). The MISTIE (Minimally Invasive Surgery plus Alteplase for Intracerebral Hemorrhage Evacuation) trials have confirmed the overall safety and improvement on mortality of MIS (Hanley, Thompson et al., 2016, Hanley, Thompson et al., 2019). During the MIS procedure, the possibility exists that rtPA may make a direct contact with the brain parenchyma around hematoma. Thus, the effect of rtPA on the surrounding tissue of the hematoma at the microscopic level needs to be explored.

rtPA, a recombinant protein of the physiological tissue-type plasminogen activator (tPA) first produced by recombinant DNA technology in 1982, dissolves the hematoma through turning fibrinogen into active fibrinolytic serine protease (Pennica, Holmes et al., 1983, Soni, Wijeratne et al., 2021). rtPA is consisted of five domains, the protease domain, the finger domain, the Epidermal Growth Factor (EGF) domain and two Kringle domains. Protease domain catalyzes the conversion of plasminogen to plasmin. The finger domain binds to endocytosis-related receptors, such as low-density lipoprotein receptor-related protein (LRP), mediating the transcytosis of rtPA by endothelial cells. The EGF domain upregulates trophic pathway through activating EGF receptor. Kringle domains play a part in plasminogen activation by stabilizing the complex of rtPA and liver clearance of rtPA(Zhu, Wan et al., 2019). It has been reported that rtPA can cause BBB damage, neuroexcitatory toxicity and inflammatory cascades (Abu Fanne, Nassar et al., 2010, Carbone, Vuilleumier et al., 2015, Kassner, Roberts et al., 2009, Lesept, Chevilley et al., 2016, Nicole, Docagne et al., 2001, Niego & Medcalf, 2014, Parcq, Bertrand et al., 2012, Pineda, Ampurdanés et al., 2012, Shi, Zou et al., 2021, Zhang, An et al., 2009). On the other hand, tPA has been shown to inhibit neuronal apoptosis, enhance synaptic plasticity and protect white matter (Correa, Gauberti et al., 2011, Haile, Wu et al., 2012, Lenoir, Varangot et al., 2019, Liang, Ding et al., 2016, Pang, Teng et al., 2004, Xia, Pu et al., 2018). The most studies on the application of rtPA have focused on ischemic stroke and less on hemorrhagic stroke, and have mostly explored aspects related to brain edema and BBB.

Therefore, the aim of this study was to investigate the direct effects and mechanisms of rtPA on neurons after experimental ICH, and clarify the specific domain that mediated the pathway activation, to provide a scientific basis for the safety of rtPA application in clinical settings.

## Results

### rtPA Attenuated Neurological Behavior Impairment After ICH

To define the effects of rtPA on neurological behavioral impairment after ICH, we performed behavioral testing before and at 6 h, 24 h, and 72 h after ICH. Compared with the sham group, there was no significant difference in the neurological function scores of mice in the sham + rtPA group (Figure 2A-C). The mice in the ICH group showed significant neurological impairment compared to the sham group, while there was no statistical difference between the ICH group and the ICH + vehicle group at 24 and 72 h after ICH (Figure 2A-C). After rtPA treatment at 12 h after ICH, the neurological impairment in ICH + rtPA group was ameliorated at 24 and 72 h after ICH compared with the ICH group and the ICH + vehicle group (Figure 2A-C). These results suggested that stereotactic injection of therapeutic doses of rtPA did not lead to neurological impairment and rtPA attenuates neurological behavior impairment after ICH.

**Figure 1.**
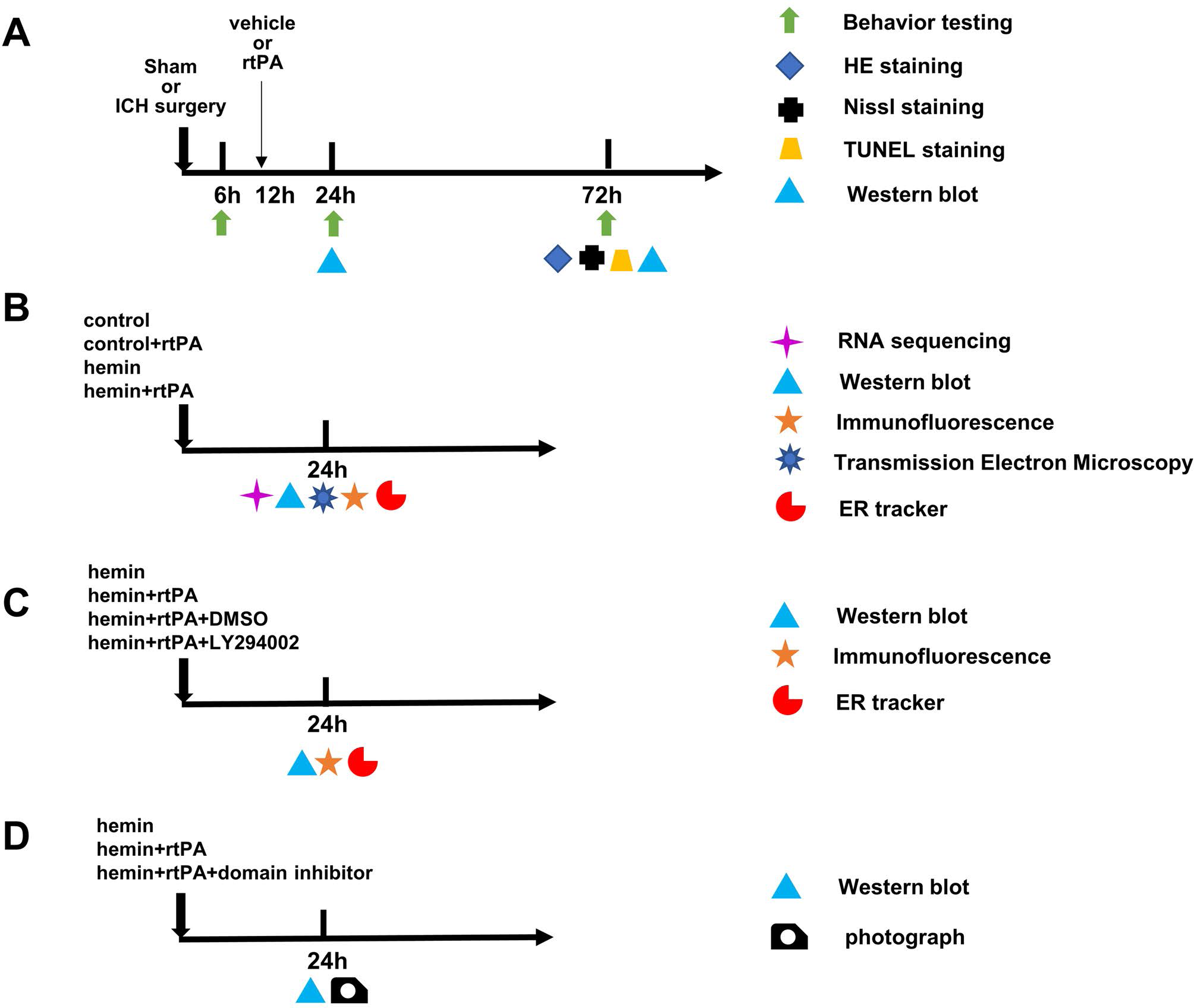
Experiment design. (A) In Experiment 1, the role of rtPA in the ICH model was explored. (B) In Experiment 2, the role of rtPA in hemorrhage stroke in vitro and the possible mechanism was explored in primary cortical neurons. (C) In Experiment 3, the mechanism of rtPA’s effect in hemorrhage stroke in vitro was explored. (D) In Experiment 4, the protein domain associated with rtPA’s neuroprotective effect in hemorrhage stroke in vitro was explored.

**Figure 2.**
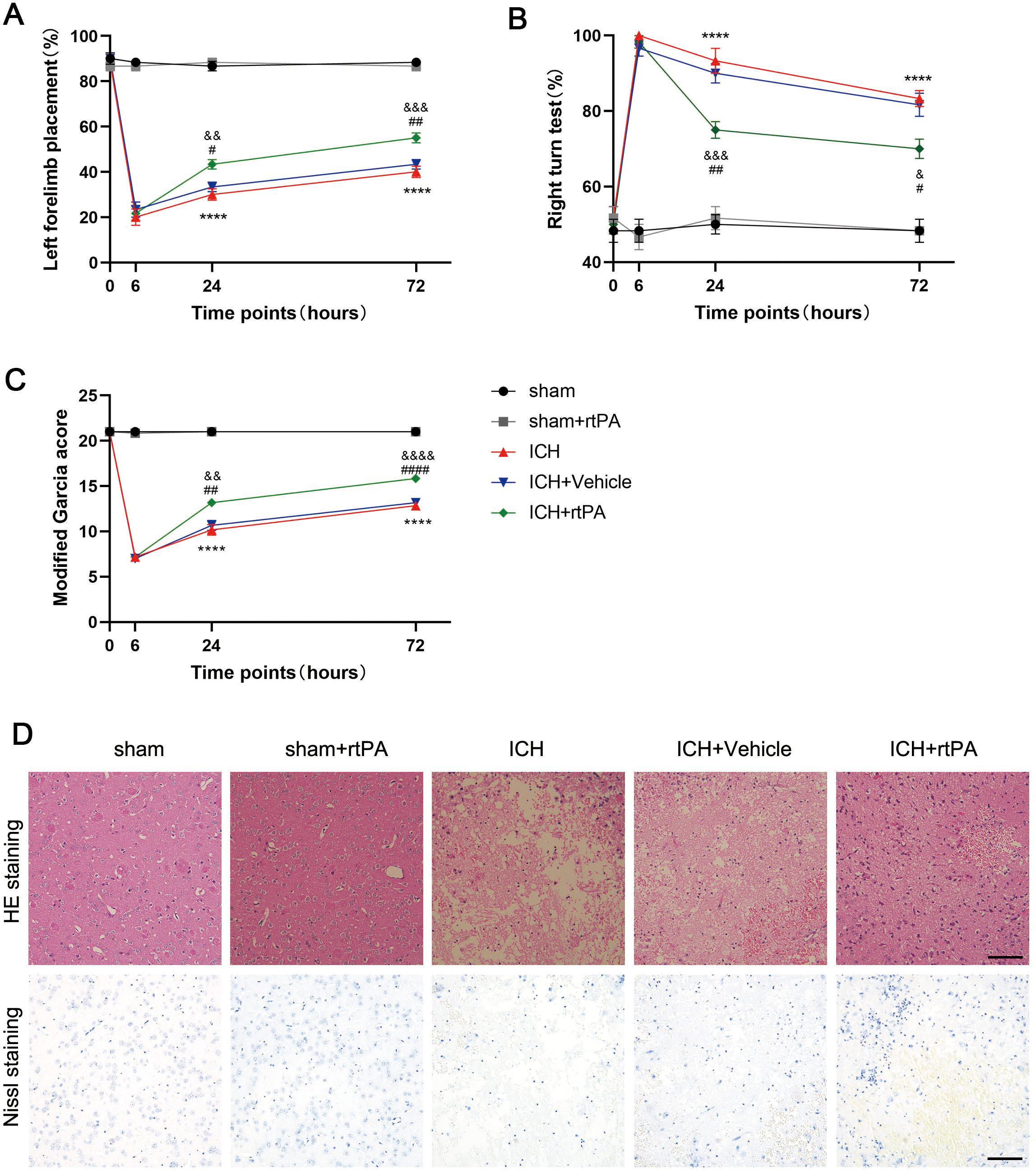
rtPA attenuated neurological behavior impairment and pathological damage after ICH. (A-C) Left forelimb placement experiment, right turn experiment, modified Garcia score conducted at 6 h, 24 h, 72 h after different treatment (n=6, \*\*\*\**P*<0.0001, compared with sham group &: *P*<0.05, &&: *P*<0.01, &&&: *P*<0.001, &&&&*P*<0.0001, compared with ICH group; #: *P*<0.05, ##*P*< 0.01, ####*P*<0.0001, compared with ICH+Vehicle group). (D) HE staining and Nissl staining at 72h after different treatment (scale bar = 100 um, n=3).

### rtPA Against the Pathological Damage at 72 h Post-ICH Induction

To explore the effects of rtPA on histopathological damage of the perihematoma after ICH, we further evaluated the pathomorphological changes of the perihematoma tissue (basal ganglia site) by HE staining and Nissl staining at 72 h after ICH. The results showed that the histomorphology of the basal ganglia region of the brain in the sham and the sham + rtPA groups did not show any significant abnormalities (Figure 2 D). Compared with the sham group, the number of neurons in the perihematoma tissue in the ICH and ICH + vehicle groups was significantly reduced and disorganized, and a large number of atrophied and deeply stained neurons were seen, while no significant difference between the ICH and ICH + vehicle groups was detected. However, rtPA treatment reduced neuropathic damage compared with ICH + vehicle group. These results suggested the therapeutic doses of rtPA did not lead to pathological damage and the rtPA attenuated the pathological damage after ICH.

### rtPA Against the Apoptosis at 24 h and 72 h After ICH

To determine the effects of rtPA in neuronal apoptosis after ICH, we examined the TUNEL staining at 72h after ICH. The TUNEL staining showed that no TUNEL-positive cells were observed in the sham and sham + rtPA groups, while a large number of apoptotic cells were observed in peri-hematoma tissue in the ICH and ICH + vehicle groups. There was no significant difference between the ICH and ICH + vehicle groups. After rtPA treatment, the number of TUNEL-positive cells was significantly reduced (Figure 3A and B).

**Figure 3.**
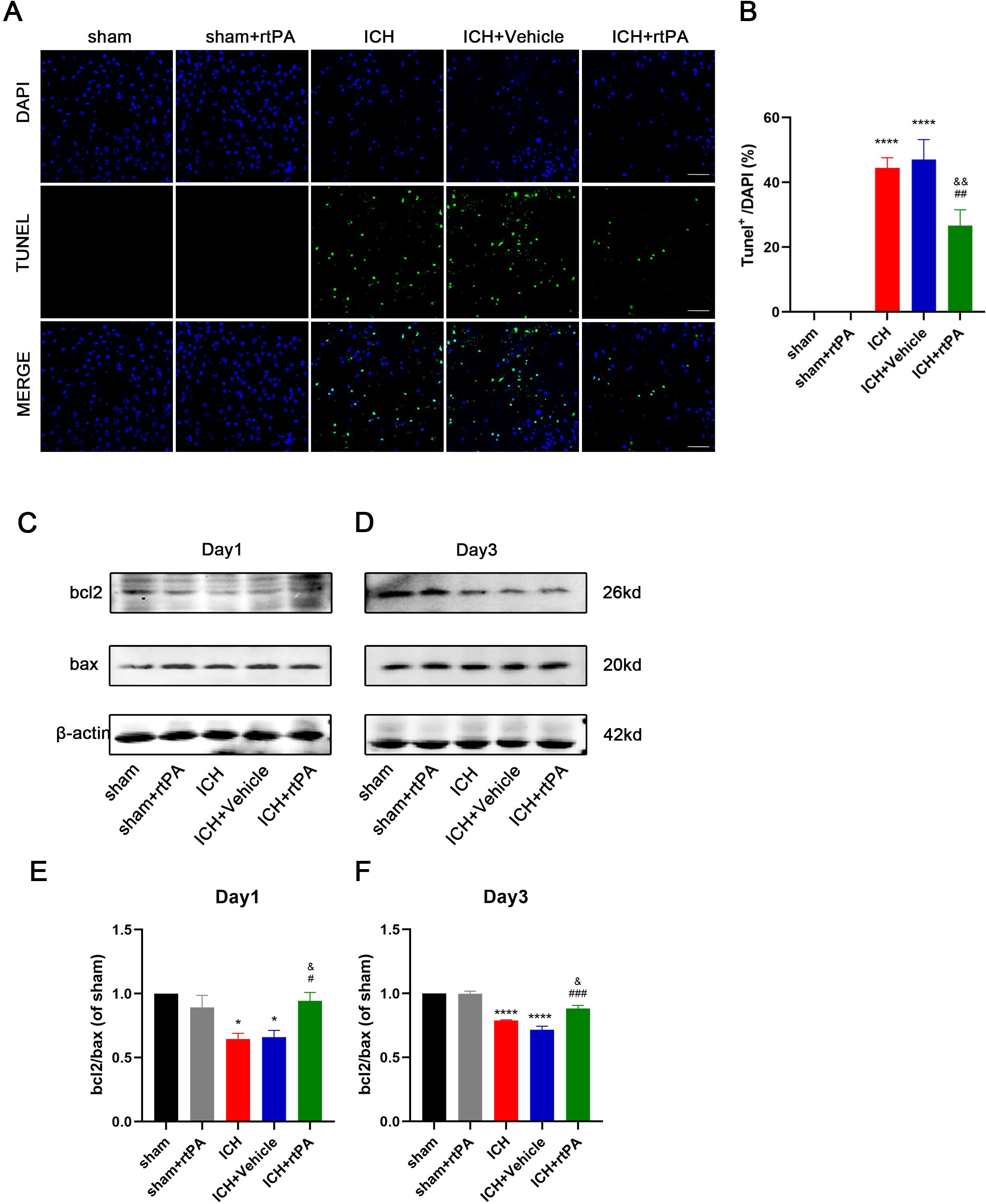
rtPA attenuated apoptosis of peri-hemotoma tissue after ICH. (A) The representative picture of Tunel staining conducted at 72 h after different treatment (scale bar = 100 um). (B) The proportion of tunel-positive cells to all nucleated cells surrounding the hematoma. (C and E) The representative band and quantitative analysis of apoptosis associated protein at 24 h after treatment. (D and F) The representative band and quantitative analysis of apoptosis associated protein at 72 h after treatment (n=3, \**P*<0.05, \*\*\*\**P*<0.0001, compared with sham group; &: *P*<0.05, &&*P*<0.01, compared with ICH group; #*P*<0.05, ##*P*<0.01, ###: *P*<0.001, compared with ICH+Vehicle group).

Additionally, we further detect the protein levels of bcl2 and bax, which are apoptosis-related proteins, by western blot analysis at 24 h and 72 h after ICH. The results showed that no significant differences were noted in the sham and sham + rtPA groups. Compared with the sham group, the anti-apoptotic factor bcl2 and the ratio of bcl2/bax decreased significantly in the ICH group and ICH + Vehicle group at 24 h and 72 h after treatment. After rtPA treatment, bcl2 expression increased, and the ratio of bcl2/bax was significantly higher compared with the ICH group and ICH + vehicle group (Figure 3C-F). These results indicated that rtPA treatment did not lead to apoptosis in normal mouse brains. rtPA is able to ameliorate neuronal apoptosis after ICH.

### rtPA Inhibited Excessive Autophagy at 24 h and 72 h After ICH

To explore the effects of rtPA in autophagy after ICH, the protein levels of beclin1, p62, and LC3-II/LC3-I, which are indicators of autophagy, were detected by western blot analysis at 24 and 72 h after ICH. The results showed that compared with the sham group, the level of beclin1 was significantly increased, the level of p62 was significantly lower, LC3-II and LC3-II/LC3-I ratios were significantly increased in the peri-hematoma tissues of the ICH and ICH + Vehicle groups at 24 h and 72 h after ICH. After administration of rtPA, the level of beclin1 was significantly decreased, the levels of p62 was significantly increased, and the ratio of LC3-II/LC3-I was decreased compared with the ICH and ICH + Vehicle groups. Meanwhile, there was no statistical difference between the sham group compared with the sham + rtPA group and between the ICH group compared with the ICH + Vehicle group (Figure 4). These results showed that rtPA did not lead to excessive autophagy in normal mouse brains. rtPA treatment can inhibit excessive autophagy after ICH.

**Figure 4.**
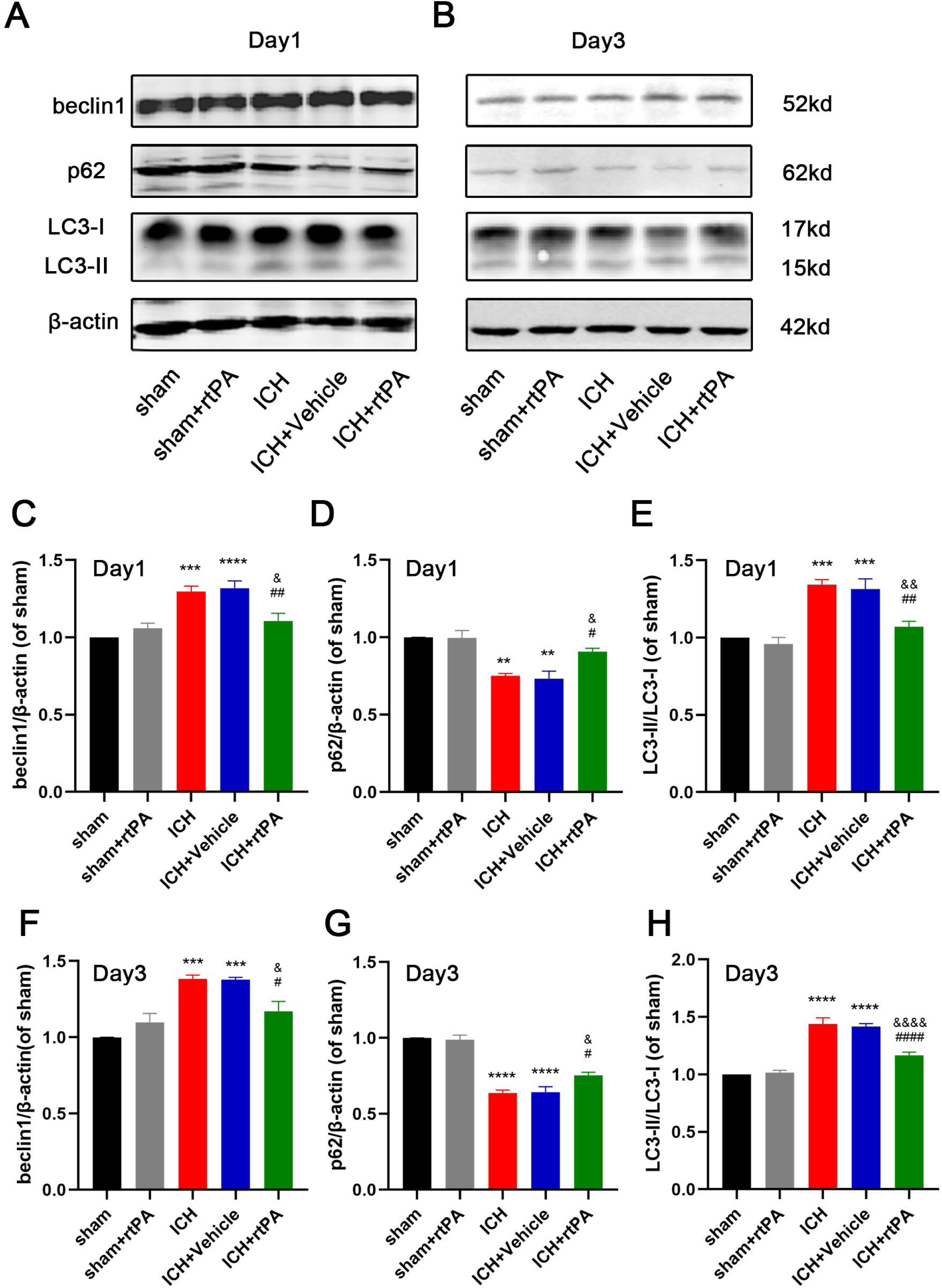
rtPA attenuated autophgy of peri-hemotoma tissue after ICH. (A, C, D, E) The representative band and quantitative analysis of autophagy associated protein at 24 h after treatment. (B, F, G, H) The representative band and quantitative analysis of autophagy associated protein at 72 h after treatment (n=3, **: *P*<0.01, ***: *P*<0.001, ****: *P*<0.0001, compared with sham group; &: *P*<0.05, &&: *P*<0.01, &&&&: *P* <0.0001, compared with ICH group; #: *P*<0.05, ##: *P*<0.01, ####: *P*<0.0001, compared with ICH+Vehicle group).

### rtPA Inhibited Apoptosis and Autophagy in Hemorrhagic Stroke In Vitro

To explore the direct effect of rtPA on neuronal apoptosis, we treated primary neuronal cells with hemin to establish the ICH model in vitro. We first explored the optimal concentrations for hemin and rtPA treatment through a Cell Counting Kit-8 (CCK-8) experiment (Fig. S1). 40 uM hemin and 300 nM rtPA were adopted in following experiment. RNA sequencing was conducted at 24 hours after drug treatment, and the heatmap of transcriptomics gene expression profile, principal component analysis and Pearson correlation analysis were shown in Fig. S2. The differential expression genes (DEGs) between control group and hemin group associated with Autophagy animal (KEGG: mmu04140), positive regulation of neuron apoptotic process (GO: 0043525), and positive regulation of response to endoplasmic reticulum stress (GO: 1905898) were screened. As shown in Figure 5, rtPA treatment reversed the expression of a mount of DEGs in these pathways. The farther western blot experiments have proved that rtPA treatment upregulated bcl2/bax ratio and p62, decreased LC3II/LC3I ratio and beclin1 (Figure 6A-B, D-G), and decreased autophagosome number detected by transmission electron microscopy (Figure 6C). These results indicated the anti-apoptosis and anti-autophagy effect of rtPA after ICH in vitro.

**Figure 5.**
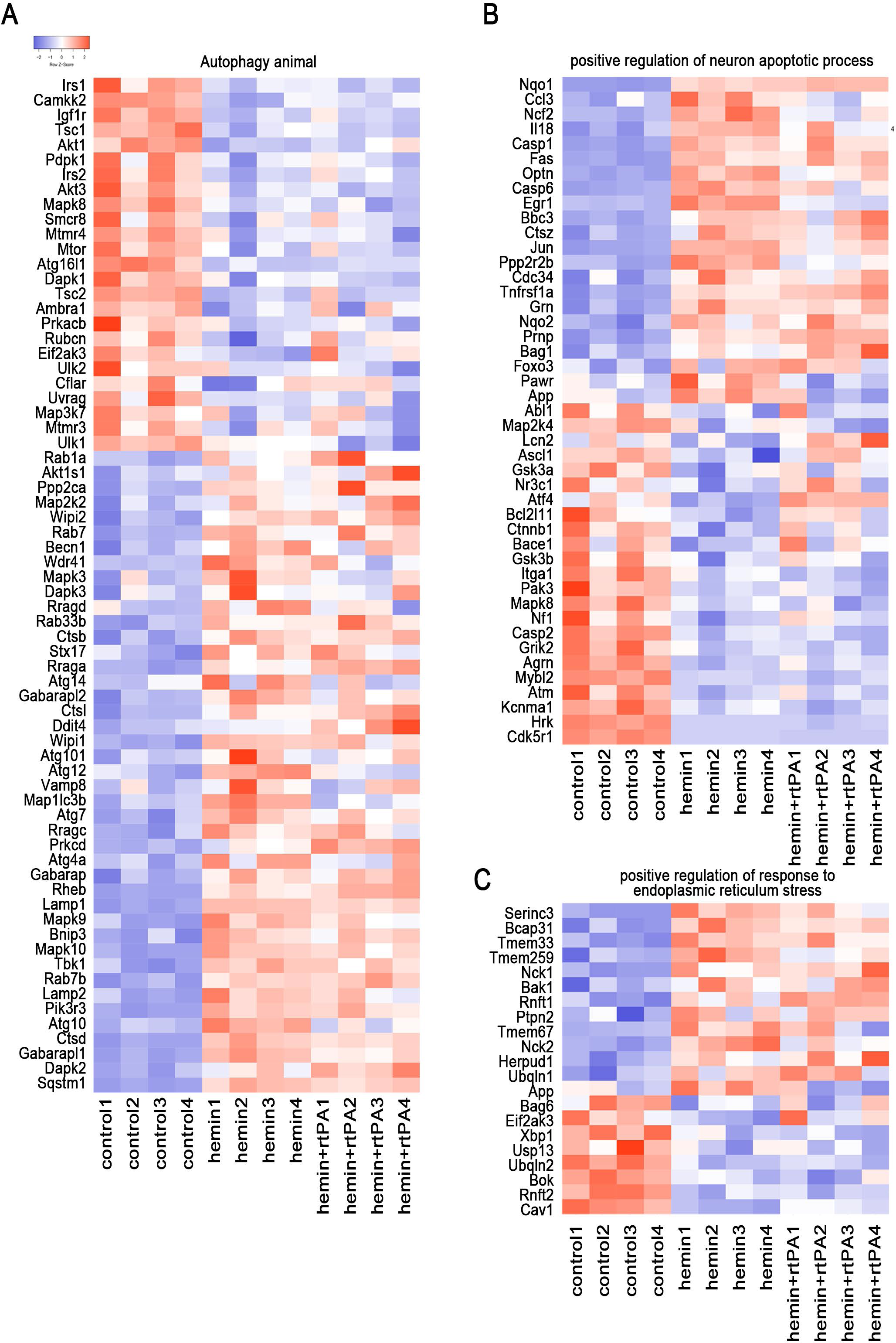
Differential Expression Genes (DEGs) on autophagy, neuron apoptosis and endoplasmic reticulum stress between different treatment. (A-C) The differential expression genes (DEGs) between control group and hemin group associated with Autophagy animal (KEGG: mmu04140), positive regulation of neuron apoptotic process (GO: 0043525), and positive regulation of response to endoplasmic reticulum stress (GO: 1905898) were screened, and the transcriptional levels of DEGs in each group were presented as heatmaps.

**Figure 6.**
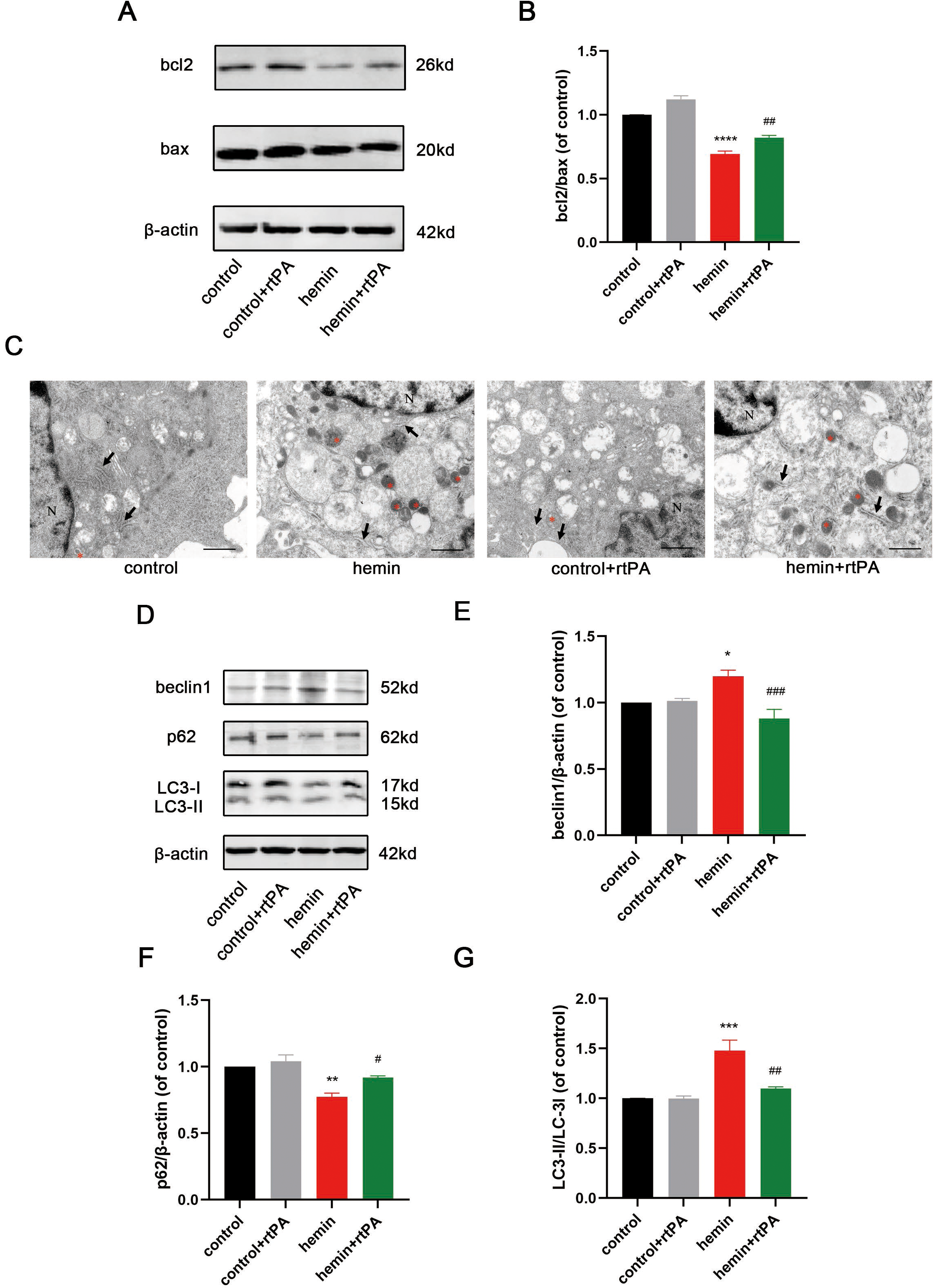
rtPA attenuated apoptosis and autophagy of neuron after experimental ICH in vitro. (A, B) The representative band and quantitative analysis of apoptosis associated protein. (C) The images of transmission electron microscopy of neuron in different treatment, with *(red) representing autophagosomes, black arrows indicating the endoplasmic reticulum, and N means nucleus. (D-G) Representative band and quantitative analysis of autophagy associated protein (n=3, *: *P*<0.05, **: *P*<0.01, ***: *P*<0.001, compared with control group; #: *P*<0.05, ##: *P*<0.01, ###: *P*< 0.001, compared with hemin group).

### rtPA Inhibited ER Stress in Hemorrhagic Stroke In Vitro

To observe the morphology and continuity of the ER in primary neurons, transmission electron microscopy was used at 24 h after hemin treatment. The results showed that in the control group and control + rtPA group, the ER was folded, normal in morphology, closely arranged, and continuous in structure At 24 h after hemin treatment, the ER lost normal morphology as reflected by swelling, severe fracture, and fragmentation, while the rtPA treatment relieve the ER damage induced by hemin. (Figure 6C).

Additionally, we used ER-tracker to label the ER at 24 h after hemin treatment and reconstructed the three-dimensional morphology of the ER by confocal microscopy. The results showed that the ER was continuous in the control and the control + rtPA groups. In hemin-neurons, the ER was severely disrupted with poor continuity, while the continuity was improved after rtPA treatment (Figure 7G).

**Figure 7.**
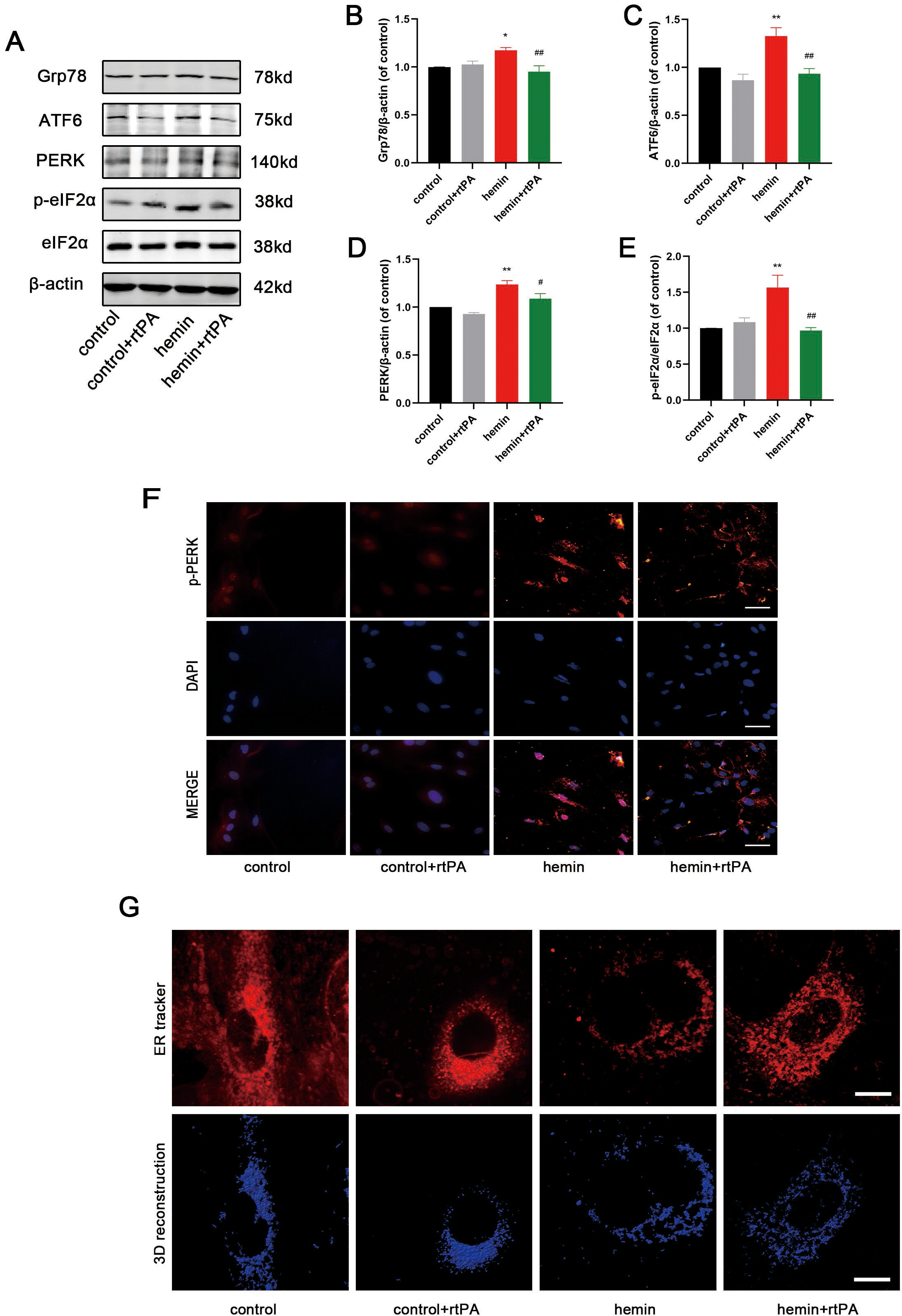
rtPA ameliorated endoplasmic reticulum stress of neuron after experimental ICH in vitro. (A-E) Representative band and quantitative analysis of ER stress associated protein (n=3, *: *P*<0.05, **: *P*<0.01, compared with control group; #: *P* <0.05, ##: *P*<0.01, compared with hemin group). (F) Immunofluorescence images of p-PERK in neurons of different groups, scale bar = 50 um. (G) Confocal images and three-dimensional reconstruction of endoplasmic reticulum continuity by ER tracker, scale bar = 3 um.

Furthermore, we detected the protein levels of Grp78, ATF6, PERK, p-eIF2α, and eIF2α, which are indicators of ER stress, by western blot analysis. The results showed that there was no significant difference in the protein levels of these indicators between the control and the control + rtPA groups. Compared with the control group, the levels of Grp78, ATF6, PERK, and p-eIF2α/eIF2α were increased in the hemin group, and rtPA treatment significantly decreased expression levels of these indicators when compared to the hemin group (Figure 7A-E).

To further define the effects of rtPA on ER stress after hemin treatment, we detected p-PERK expression by immunofluorescence in primary neurons at 24 h after hemin treatment. The results showed that the expression of p-PERK was barely observed in the control and control + rtPA groups, while hemin treatment caused an increase in the fluorescence intensity of p-PERK, which was reduced after rtPA treatment (Figure 7F). These results showed that rtPA had no effect on ER stress in normal neurons while neuronal ER stress levels were increased in hemorrhagic stroke in vitro. rtPA treatment can alleviate hemin-induced ER stress in vitro.

### rtPA Inhibited neuron Apoptosis, Excessive Autophagy, and ER Stress by Upregulating PI3K/AKT/mTOR in Hemorrhagic Stroke In Vitro

To further explore the mechanism of the direct protective effect of rtPA on neurons, we found that the expression profiles of PI3K/AKT and mTOR pathway changed significantly after hemin treatment, and the hemin + rtPA group had a trend of gene expression shifting toward the control group (Figure 8A and B). The results indicated that rtPA had a certain regulatory effect on PI3K/AKT pathway and mTOR pathway after hemin treatment, and the direct protective effect of rtPA may be related to this pathway.

**Figure 8.**
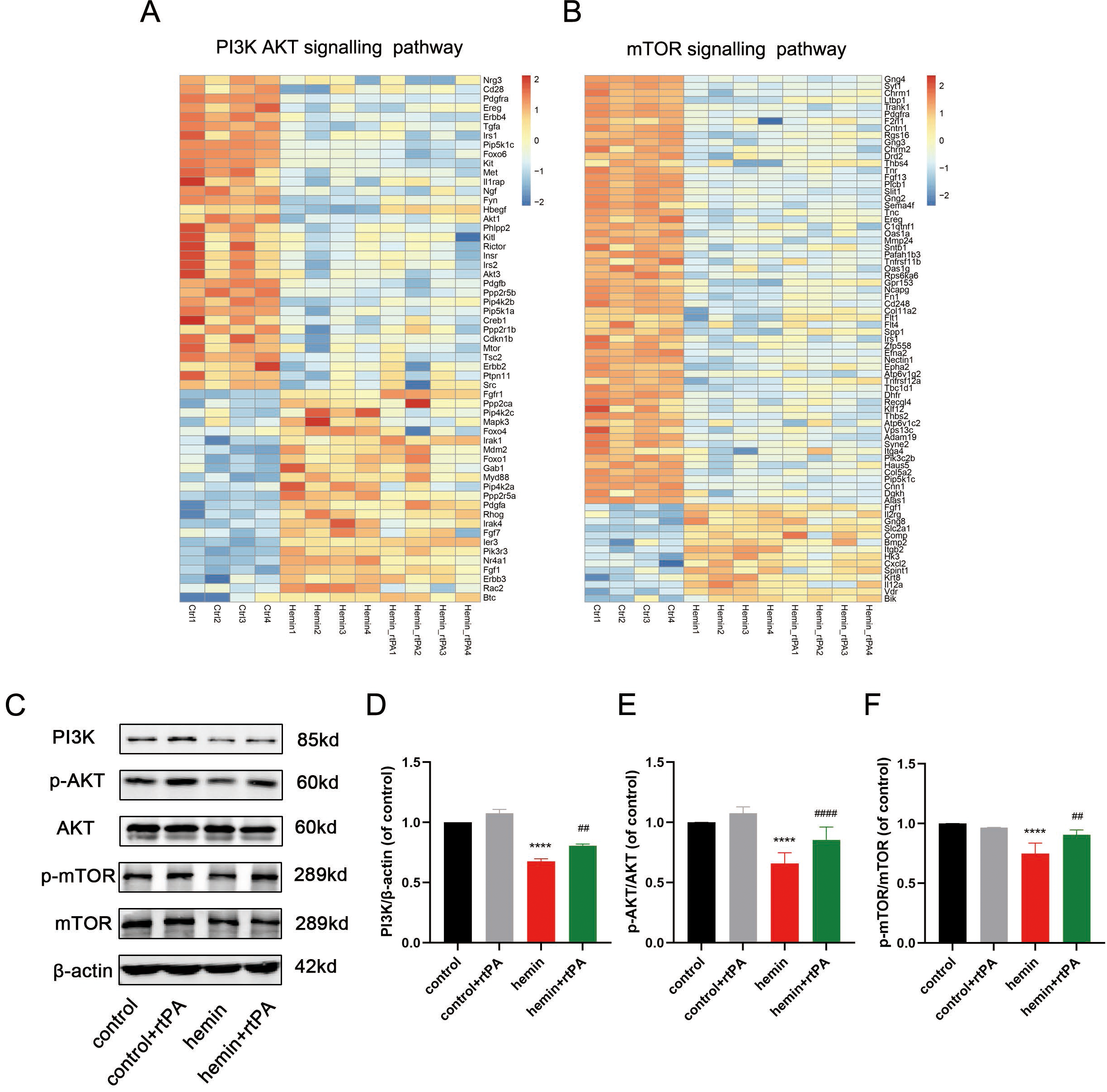
rtPA regulated PI3K-AKT-mTOR pathway of neuron after experimental ICH in vitro. (A, B) The differential expression genes (DEGs) between control group and hemin group associated with PI3K/AKTpathway (KEGG : mmu04151) and mTOR pathway (KEGG: mmu04150) were screened, and the transcriptional levels of DEGs in each group were presented as heatmaps. (C-F) Representative band and quantitative analysis of PI3K-p85, p-AKT, AKT, P-mTOR, mTOR (n=3, ****: *P*< 0.0001, compared with control group; ##: *P*<0.01, ####: *P*<0.0001compared with hemin group).

Thus, we detected the changes in protein levels of PI3K/AKT/mTOR pathway proteins, including PI3K p85, p-AKT, AKT, p-mTOR, and mTOR, by western blot analysis at 24 h after hemin treatment. The results showed that there was no statistical difference in levels of PI3K p85, p-AKT/AKT and p-mTOR/mTOR between the control and control + rtPA groups, indicating that rtPA had no effect on the PI3K/AKT/mTOR signaling pathway in normal neurons. After hemin treatment, levels of PI3K p85, p-AKT/AKT, and p-mTOR/mTOR were significantly downregulated compared to the control group. However, the rtPA treatment upregulated the levels of PI3K p85, p-AKT/AKT, and p-mTOR/mTOR (Figure 8C-F). Next, we explored whether the PI3K pathway inhibitor LY294002 could reverse the neuroprotective effects of rtPA in hemorrhagic stroke in vitro. We found that LY294002 could effectively reverse the upregulation of levels of PI3K p85, p-

AKT/AKT, and p-mTOR/mTOR by rtPA in hemin-neurons using western blot analysis (Figure 9A-D). We further investigated whether LY294002 could reverse the anti-apoptosis, anti-autophagy, and anti-ER stress effects of rtPA on primary neurons after hemin treatment. Western blotting demonstrated that LY294002 treatment reversed the regulatory effects of rtPA on apoptosis-related proteins (bcl2 and bax), autophagy-related proteins (beclin1, p62, and LC3-II/LC3-I), and ER stress-related proteins (ATF6, p-eIF2α, and eIF2α) (Figure 9E-J, 10C-E). ER tracker and 3D reconstruction of ER showed that the effect of rtPA to improve ER continuity after hemin treatment was attenuated by LY294002 (Figure 10A). LY294002 treatment reversed the downregulation of fluorescence intensity of p-PERK induced by rtPA (Figure 10B). These results showed that the treatment of PI3K inhibitor LY294002 reversed the protective effects of rtPA in apoptosis, autophagy, and ER stress in hemorrhagic stroke in vitro.

**Figure 9.**
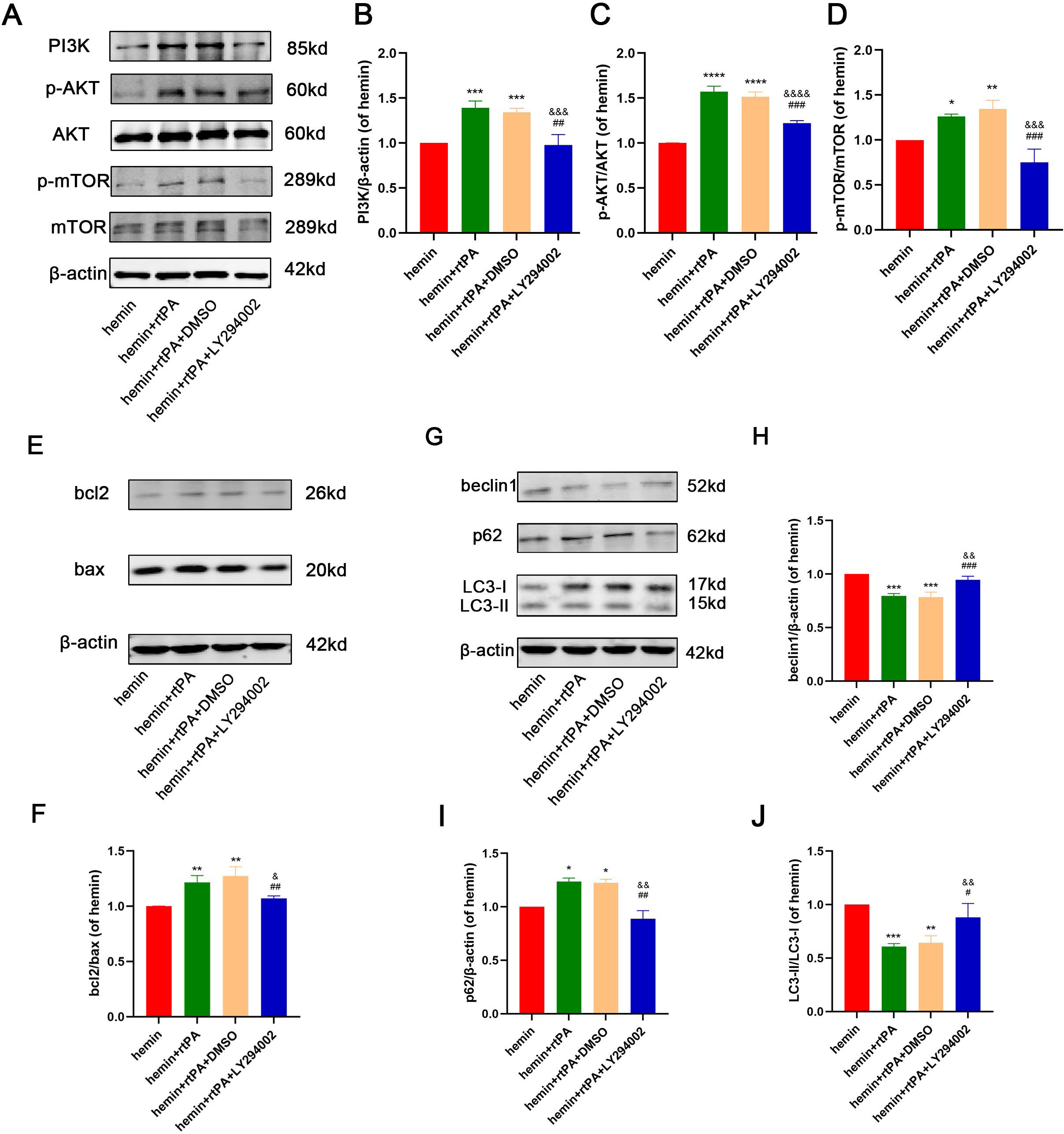
PI3K inhibitor LY294002 reversed the anti-apoptosis and anti-autophagy effect of rtPA. (A-D) Representative band and quantitative analysis of PI3K-p85, p-AKT, AKT, P-mTOR, mTOR. (E, F) Representative band and quantitative analysis of apoptosis associated protein. (G-J) Representative band and quantitative analysis of autophagy associated protein (n=3, *: *P*<0.05, **: *P*<0.01, ***: *P*<0.001, ****: *P* <0.0001, compared with hemin group; &: *P*<0.05, &&: *P*<0.01, &&&: *P*<0.001, &&&&: *P*<0.0001, compared with hemin + rtPA group; #: *P*<0.05, ##: *P*<0.01, ###: *P*<0.001, compared with hemin + rtPA + DMSO group).

**Figure 10.**
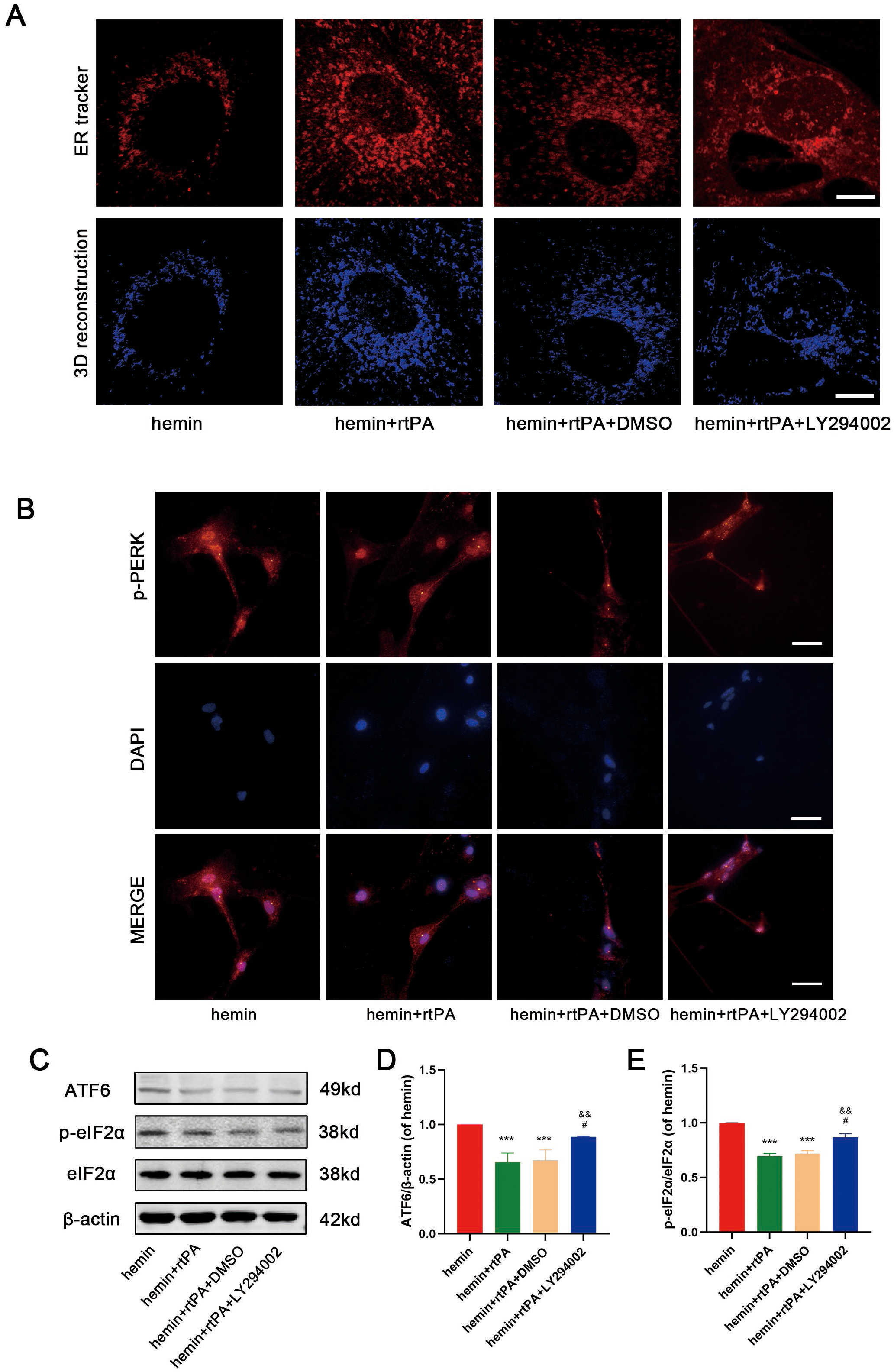
PI3K inhibitor LY294002 reverse the anti-ER stress effect of rtPA. (A) Confocal images and three-dimensional reconstruction of endoplasmic reticulum continuity by ER tracker, scale bar = 3 um. (B) Immunofluorescence images of p-PERK in neurons of different groups, scale bar = 50 um. (C-E) Representative band and quantitative analysis of ER stress associated protein (n=3, ***: *P* < 0.001, compared with hemin group; &&: *P*<0.01, compared with hemin + rtPA group; #: *P* <0.05, compared with hemin + rtPA + DMSO group).

### EGF Domain of rtPA Might Upregulated PI3K/AKT pathway in Hemorrhagic Stroke In Vitro

In order to further explore the specific domain of rtPA exerting neuroprotective effects, we used different inhibitors to inhibit rtPA domains’ effect, GGACK for the protease domain, AG1487 for the EGF domain, RAP for the finger domain, TXA for the Kringle domain. The result showed that AG1487 reversed the neuroprotective effect of rtPA, with neurons in hemin + rtPA+ AG1487 group showing obvious neural damage observed by microscopy (Figure 11D). Meanwhile, AG1487 significantly downregulated the expression level of p-AKT and somewhat reduced the expression level of PI3K p85 (but no statistical difference) (Figure 11A-C).

**Figure 11.**
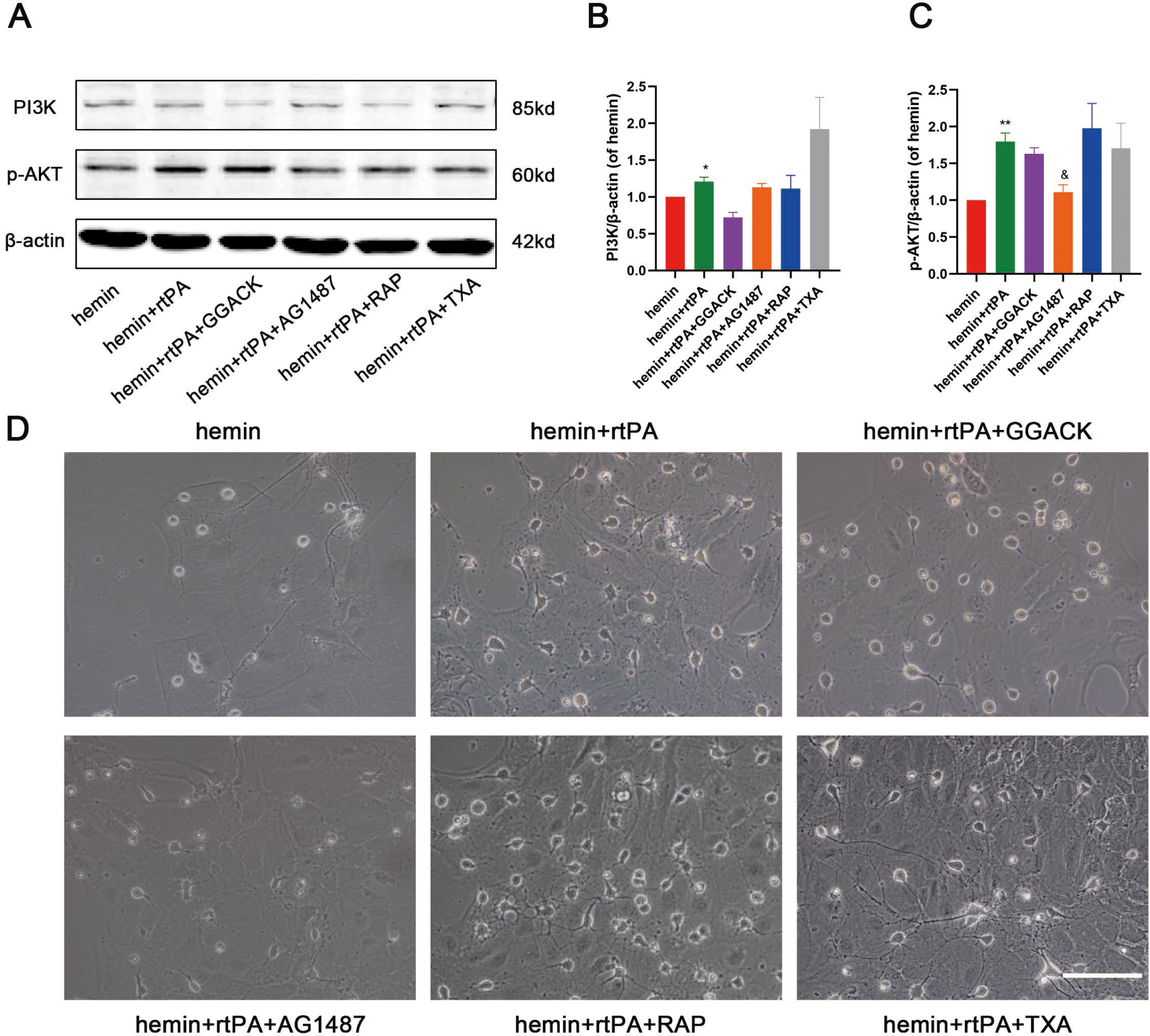
EGF Domain of rtPA Might Upregulated PI3K/AKT pathway in Hemorrhagic Stroke In Vitro. (A-C) Representative band and quantitative analysis of PI3K p85, p-AKT (n=3, *: *P*<0.05, **: *P*<0.01, campared with hemin group; &: *P* <0.05, campared with hemin + rtPA group). (D) The microscopic images of different treatment, scale bar = 100 um.

## Discussion

The mechanisms of neurological impairment after ICH are complex. The compression of the surrounding brain tissue by the hematoma after ICH and the processes of blood-brain barrier damage and brain edema induced by toxic substances from hematoma are the main pathophysiological mechanisms of neurological impairment. The minimally invasive surgery plus rtPA is a common used hematoma removal technique for ICH. However, this therapy arouses a question whether the direct contact of rtPA with brain tissue cause neurological damage. In this study, we found that stereotactic injection rtPA without hematoma aspiration significantly reduced ICH-induced neurological impairment, histopathological damage, apoptosis and excessive autophagy without affecting the neurological function of normal mice. This finding suggested the rtPA was likely to have a direct neuroprotective effect in addition to the hematoma-liquefaction ability. However, the neuroprotective effect of rtPA may be caused by the hematoma-liquefaction ability of rtPA, which reduces the hematoma volume and subsequently reduces the direct and indirect damage caused by the hematoma. Therefore, we further explored the direct effect of rtPA on neurons in hemin-treated primary neurons. We found that rtPA could inhibited apoptosis, excessive autophagy and ER stress in hemorrhagic stroke in vitro through PI3K-AKT-mTOR pathway.

Apoptosis was the main mechanism of early injury perihematoma tissue after ICH (Li, Han et al., 2017, Matsushita, Meng et al., 2000, Zille, Karuppagounder et al., 2017). Inhibition of apoptosis is extremely important for cell survival after ICH (Li, Song et al., 2021, Liu, Liu et al., 2021). BAX and Bcl-2 are important genes that regulate apoptosis (Oltvai, Milliman et al., 1993). In short, BAX promotes apoptosis, and Bcl-2 inhibits apoptosis, thus the ratio of bcl-2/ bax can reflect whether the cells tend to undergo apoptosis or survive after stimulation. In this study, we found that the ratio of bcl-2/ bax in hemin-treated neurons decreased in vitro, but this decrease was alleviated by rtPA treatment. This study showed that rtPA can directly reduce apoptosis in hemorrhagic stroke in vitro.

Autophagy is closely linked to apoptosis. Autophagy has a dual role in the ICH. The moderate autophagy contributes to cell survival, whereas excessive autophagy can induce apoptotic processes or even lead to cell death (Ni, Wei et al., 2022). In a rat model of cerebral hemorrhage, autophagy and endoplasmic reticulum stress are enhanced within one week after ICH. The excessive autophagy is involved in endoplasmic reticulum stress-induced brain injury within 6 h after ICH, and at 7 d after cerebral hemorrhage, autophagy can play a protective role by removing damaged cellular debris (Duan, Wang et al., 2017). Excessive endoplasmic reticulum stress can induce autophagy through multiple pathways, such as the PERK-elF2α and IRE-1α pathways, and can also upregulate the expression of pro-apoptotic signals of caspase-3 to induce apoptosis (Cao, Peng et al., 2021). Inhibition of the classic pathway of endoplasmic reticulum stress (Grp78-IRE1/PERK) reduces apoptosis in a rat model of cerebral hemorrhage (Huang, Lan et al., 2018). In experimental brain hemorrhage models, the release of toxic substances within the hematoma, such as iron, activates the autophagic process (He, Wan et al., 2008), and the use of iron-regulating agents can reduce the iron content in brain tissue and thus reduce autophagy after cerebral hemorrhage in rats (Zhang, Qian et al., 2021). The use of drugs to inhibit autophagy levels in brain hemorrhage has been shown to reduce neuronal damage (Wu, Zou et al., 2016, Zhao, Gao et al., 2021). Inhibition of the MST4/AKT signaling pathway can also improve neurological function in mice by reducing the level of autophagy after cerebral hemorrhage (Wu, Wu et al., 2020). We found that rtPA had no effect on apoptosis, autophagy, and endoplasmic reticulum stress levels in normal neurons; whereas in an in vitro brain hemorrhage model, neuronal apoptosis, autophagy, and endoplasmic reticulum stress levels were increased, and rtPA treatment was able to downregulate apoptosis levels after hemin stimulation, inhibit excessive autophagy, and attenuate endoplasmic reticulum stress.

As an endogenous factor synthesized by the nervous system, tPA plays an important role in cell growth and migration in the nervous system. The role of tPA in the nervous system still needs to be further explored. On the one hand, tPA may aggravate injury by disrupting the blood-brain barrier, mediating excitotoxic effects, and amplifying inflammatory effects. On the other hand, tPA may play a protective role in promoting neuronal survival, facilitating synaptic remodeling, and protecting white matter. Exogenously administered rtPA also has effects on the homeostasis and function of the nervous system. In clinical applications in the ischemic stroke, rtPA has the disadvantage of a short time window, which can aggravate BBB destruction and neuroinflammation, but it also has the effects of protecting neurons, promoting synaptic remodeling, and improving white matter damage. Numerous studies have shown that tPA may have potential for neuroprotective effects in addition to thrombolytic effects in CNS diseases. In experimental ischemic models, tPA can increase IGF1 serum levels through protein hydrolysis, thereby inhibiting excessive autophagy and attenuating cell death (Thiebaut, Buendia et al., 2021). In ischemia-reperfusion (IR), tPA regulates the mitochondrial autophagic process by increasing AMPK phosphorylation and FUNDC1 expression, thereby inhibiting apoptosis and protecting neurons (Cai, Yang et al., 2021). In a rat model of traumatic brain injury, inhibition of endogenous tPA exacerbates neuronal apoptosis and axonal injury (Zhao, Liu et al., 2020). In the in vitro oxygen and glucose deprivation (OGD) model, stressed neurons release endogenous tPA, which attenuates neuronal apoptosis via the EGF-EGFR pathway, and administration of exogenous tPA to tPA knockout mice also reduces neuronal apoptosis, thus providing neuroprotection (Lemarchand, Maubert et al., 2016). In experimental ischemic stroke mice, tPA cleaves IGFBP3 to increase IGF1 plasma levels and induce IGFR activation, thereby attenuating autophagy after cerebral ischemia (Thiebaut et al., 2021).

Most of the basic studies on the application of rtPA in cerebral hemorrhage have focused on its causing brain edema and damaging the Blood-brain Barrier (BBB) (Keric, Maier et al., 2012, Rohde, Rohde et al., 2002, Tan, Chen et al., 2017). Intracerebroventricular injection of tPA leads to increased BBB permeability in normal mice (Yepes, Sandkvist et al., 2003). In a porcine model of cerebral hemorrhage, rtPA application exacerbates delayed brain edema, a process that can be blocked by fibrinogen activation inhibitors (Keric et al., 2012). In contrast, retrospective clinical studies concluded that administration of rtPA to treat patients with cerebral hemorrhage complicated by intraventricular hemorrhage did not result in increased perihematomal edema (PHE) (Volbers, Wagner et al., 2013), and other clinical studies have obtained similar conclusions (Lian, Xu et al., 2014, Ziai, Moullaali et al., 2013). It has been shown that rtPA does not exacerbate the extent of Evans blue dye extravasation after cerebral hemorrhage in mice (Pfeilschifter, Kashefiolasl et al., 2013). In a rat model of cerebral hemorrhage, rtPA application attenuated the degree of Evans blue dye extravasation after cerebral hemorrhage, but it upregulated matrix metalloproteinase (MMP)-9 expression and agonized the NF-κB pathway (Tan et al., 2017). Activated MMPs degrade the tight junction proteins of the BBB, leading to increased BBB permeability (Yang & Rosenberg, 2011). However, we focus on the survival status of perihematoma neurons and explored the role of rtPA in reducing apoptosis and autophagy levels in an animal model of ICH. We found that rtPA downregulated apoptosis and autophagy levels in perihematomal tissues of ICH mice during the acute and subacute phases, reduced perihematomal histopathological damage, thus protected the neurons. Moreover, we found that rtPA inhibited apoptosis, excessive autophagy and ER stress in hemorrhagic stroke in vitro.

The PI3K/AKT/mTOR pathway mediates many processes of cell growth, which are also closely related to autophagy regulation (Jadaun, Sharma et al., 2021, Porta, Paglino et al., 2014). Decreased mTOR expression and increased autophagy levels in perihematoma tissues were detected in a mouse model of cerebral hemorrhage (Yu, Zhang et al., 2017). tPA can promote PI3K/AKT/mTOR pathway activation through multiple pathways. In vitro experiments by Grummisch et al. revealed that tPA promotes the survival of primary neurons through an mTOR-dependent mechanism (Grummisch, Jadavji et al., 2016). Moreover, tPA can interact with LDL-related protein-1 (LRP-1) through the finger-like structural domain, induce dose-dependent migration of neutrophils through the LRP-1/AKT pathway, and contribute to granulocyte degranulation through pathways such as PI3K/AKT and ERK1/2 (Liberale, Bertolotto et al., 2020). In a serum deprivation model, tPA exerts anti-apoptotic effects through upregulation of the PI3K pathway (Liot, Roussel et al., 2006). In an ischemic stroke model, tPA was found to increase serum levels of Insulin Growth Factor 1 (IGF1) and agonize IGF Receptor (IGFR) to regulate the PI3K/AKT/mTOR pathway through its protein hydrolysis effect, thereby reducing autophagy, attenuating cell death, and mediating neuroprotection (Thiebaut et al., 2021). In our study, we applied the PI3K inhibitor LY294002, and found that the treatment of PI3K inhibitor reversed the protective effects of rtPA in apoptosis, autophagy, and ER stress in hemorrhagic stroke in vitro. The results suggested that the protective effects of rtPA can reduce apoptosis, inhibit excessive autophagy, and inhibit ER stress by PI3K/AKT/mTOR pathway in hemorrhagic stroke in vitro. And the EGF domain might be responsible for the regulation of PI3K/AKT/mTOR pathway of rtPA.

One of the limitations of the present study is that the direct protective effect of rtPA on neurons and the molecular mechanism only were explored in hemorrhagic stroke in vitro, and an in vivo ICH model was lacking. Therefore, the mechanism related to the direct protective effect of rtPA on neurons in vivo remains unclear. Furthermore, no ER-stress agonist/inhibitor was applied, so whether the effect of rtPA in attenuating excessive autophagy and apoptosis through inhibiting ER-stress remains unclear.

In conclusion, we found that rtPA can directly protect neurons after ICH both in vivo and in vitro. This neuroprotective effect may be induced by inhibiting apoptosis, excessive autophagy, and ER Stress through PI3K/AKT/mTOR Pathway. Our study provides a rationale for investigating the safety and efficacy of rtPA in ICH (Figure 12).

**Figure 12.**
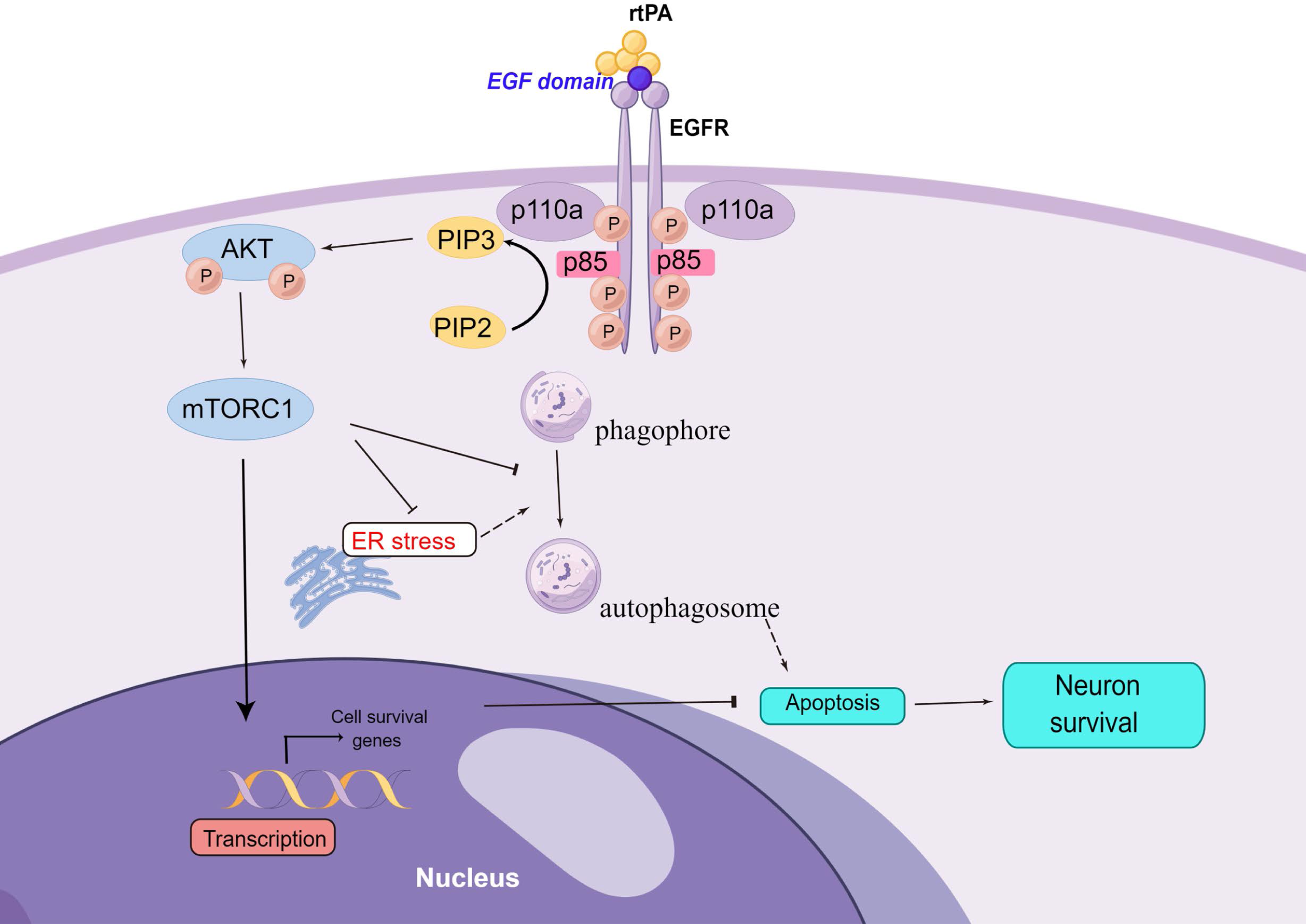
EGF domain of rtPA directly protects neurons after intracerebral hemorrhage by inhibiting apoptosis, excessive autophagy, and ER stress through PI3K/AKT/mTOR pathway.

## Materials and Methods

### Experimental Design

The experimental design is shown in Figure 1.

In Experiment 1, the role of rtPA in the ICH model was explored. Adult male C57BL/6J mice weighing 20-25 g (provided by Experimental Animal Center of Tongji Medical College, Huazhong University of Science and Technology, Wuhan, China) were randomly and equally divided into the following five groups: sham group, sham + rtPA group, ICH group, ICH + vehicle group, and ICH + rtPA group (n=6 in each group), using the table of random numbers by a technician who did not take part in this research. Behavioral testing was performed before and at 6 h, 24 h, and 72 h after ICH. Mice were sacrificed for western blotting, HE staining, Nissl staining and TUNEL staining after behavioral testing at 72h post-ICH (Figure 1A).

In Experiment 2, the role of rtPA in hemorrhage stroke in vitro and the possible mechanism was explored in primary cortical neurons. The neurons were then divided into four groups for RNA sequencing analysis, western blotting, electron microscopy, ER-tracker and immunofluorescence at 24 h after drug treatment, as follows: control group; control + rtPA group was given rtPA alone; hemin group was given hemin to induce hemorrhage stroke in vitro; hemin + rtPA group was given rtPA treatment was given at the same time as hemin (Figure 1B).

In Experiment 3, the mechanism of rtPA’s effect in hemorrhage stroke in vitro was explored. The primary cortical neurons were then divided into four groups for western blotting, ER-tracker and immunofluorescence at 24 h after drug treatment, as follows: hemin group; hemin + rtPA group; hemin + rtPA+ DMSO group; hemin + rtPA + LY294002 (PI3K inhibitor) group (Figure 1C).

In Experiment 4, the domain associated with rtPA’s neuroprotective effect in hemorrhage stroke in vitro was explored by western blot. Different inhibitor disolved in culture medium was applied to block the effect of different domain, 300 nM GGACK to block protease domain, 500 nM receptor-associated protein (RAP) (inhibitor of LRP) to block finger domain’s effect, 5 uM AG-1487 (HY13524-10, MCE) (inhibitor of EGF receptor) to block EGF domain’s effect, 300 nM tranexamic acid (TXA) (HY-B0149-5, MCE) to block a target on Kringle domain (Figure 1D).

### Animals

C57BL/6J mice were provided by the Experimental Animal Center of Tongji Medical College, Huazhong University of Science and Technology. 2-3 months-old male mice, weighing 20-25 g were adopted in our experiment. Mice were housed at 68°F to 72°F, 30% to 70% humidity in the IVC animal room under a 12-hour light/dark cycle, with food and water freely accessible.

### ICH Procedure and Treatment Strategy

An in vivo ICH model was induced via collagenase Ⅶ injection into right basal ganglia in C57BL/6J mice. After mice were anesthetized, a mini hole was drilled on skull (stereotaxic coordinates: 0.5 mm posterior to the bregma, 2.0 mm right of the middle line and 3.5 mm deep from the skull surface). 1 ul Collagenase Ⅶ (0.1 U in PBS, Sigma-Aldrich) or 1 ul PBS were infused into the right basal ganglia at a rate of 0.2 μL/min. At 12 h after ICH induction, 1 ul rtPA (1 μg in saline) or 1 ul saline was infused in the same manner.

### Behavioral Testing

Behavioral testings were performed before and at 6 h, 24 h, and 72 h after ICH. All the mice in each group were examined using left forelimb placement test, right turn test, and modified Garcia score. Mice with ICH surgery that did not exhibit obvious neurological defects were excluded from this study.

### H&E staining

Specimens were perfused with saline to remove blood cells and transversally sectioned (4-μm thickness). After fixing in 4% paraformaldehyde for 24 h, the specimens were embedded in paraffin wax. Then, the sections were used for H&E staining and imaged under a light microscope to evaluate histological changes.

### Nissl Staining

After coronal sections had been deparaffinized and rehydrated, the slides were stained in toluidine blue solution (Servicebio) for 5 min at room temperature. After clearing in distilled water, the slides were gradually dehydrated for 3 min in successive baths of ethanol, with one pass each in 70, 80, and 95% and two passes in 100%. All slides were then given two 5 min passes in 100% dimethylbenzene and coverslips were applied with neutral balsam. Finally, the numbers of surviving neurons per 400× field within the basal ganglion area were counted.

### TUNEL Staining

Cell apoptosis of brain tissues around the hematoma was measured with a TUNEL Apoptosis Assay Kit (Roche, 11684817910) according to the manufacturer’s instructions. The TUNEL-positive cells in the brain tissues around the hematoma were observed using a fluorescence microscope (Olympus BX53) and TUNEL-positive cell numbers per field were counted by another person.

### Western Blot Analysis

Brain tissue around the hematoma was separated for western blot analysis. The brain samples were washed in ice-cold PBS to remove blood, then were transferred to filter paper, carefully separated the tissue around the hematoma and weighed. Protein extraction and concentration determination were conducted as described previously(Wang, Li et al., 2021). The protein samples (20 μg/lane) were separated by 8, 10 or 12% SDS polyacrylamide gel and transferred to polyvinylidene fluoride membranes (Thermo Scientific, USA). The membranes were blocked with 5% skim milk for 1 h at room temperature and then incubated with primary antibodies overnight at 4°C. The membranes were then washed with TBST and incubated with DyLight™ 800 4X PEG-conjugated or DyLight™ 680 4X PEG-conjugated secondary antibody for 2 h at room temperature. The protein bands were visualized using Odyssey two-color infrared laser system (LICOR, USA), and the relative protein quantity was determined using ImageJ software (National Institutes of Health, USA). For the western blot analysis in cell culture of primary cortical neurons, the cell cultures were washed 2-3 times with ice-cold PBS. Then added 50 ul of cell lysate (the same as tissue homogenate), transferred to a 1.5 ml EP tube. The rest steps were the same as the western blot analysis in animal tissue proteins. The primary antibodies used include the following: Bcl-2 (Proteintech, 12789-1-AP, 1:1000), BAX (Proteintech, 50599-2-Ig, 1:1000), Beclin-1 (Proteintech, 11306-1-AP, 1:1000), SQSTM1/p62 (Abclonal, A11250, 1:1000), LC3B (Abcam, ab48394, 1:1000), GRP78/BIP (Proteintech, 66574-1-Ig, 1:1000), ATF6 (Proteintech, 24169-1-AP, 1:1000), PERK (CST, 3192S, 1:1000), Phospho-PERK (CST, 3179S, 1:1000), eIF2α (CST, 9722S, 1:1000), Phospho-eIF2α (CST, 9721S, 1:1000), PI3 Kinase p85 (CST, 4257S, 1:1000), AKT (CST, 4691S, 1:1000), Phospho-AKT (CST, 4060S, 1:1000), mTOR (CST, 2983S, 1:1000), Phospho-mTOR (CST, 2971S, 1:1000) and β-actin (Proteintech, 66009-1-Ig, 1:1000) as a loading control.

### Cell Culture of Primary Cortical Neuron

Cortical tissues were isolated from neonatal C57/BL6 mouse brains. After cutting the scalp and skull, the whole brain was removed to cold DMEM-high glucose (DMEM-HG) medium (HyClone, SH30234.01). After meninges and blood vessels were carefully peeled off, cerebral cortex was isolated and transferred to a glass dish with DMEM-HG. The cortical tissue was shreded and pipetted 20 times slightly using the pipette to disperse. The cell suspension was transferred to 2 mg/ml papain (Worthington, USA) at a volume ratio of 1:1 with DMEM-HG. After 20 min digestion, cells were filtered with a 40um cell stainer (BD, USA). The filtrate was collected in a new centrifuge tube and centrifuged at 1000 r/m at 4°C for 5 min. After discarding the supernatant, cells were resuspended with 1 ml DMEM/F12 medium (Gibco,) containing 20% FBS (Celligent, CG0430B). Cells were inoculated into 96-well plate for CCK8 experiment, 24-well plate for immunofluorescence and 6-well plate for western blot. All plates were pre-coated with poly-L-lysine hydrobromide (Sigma Aldrich, P2636). 3 h later, the medium was changed with freshly prepared primary neuronal complete culture medium composed of Neurobasal, 2% B27 and 1% penicillin-streptomycin. Cells were cultured for 14 days before the following treatment, with half of the media exchanged for fresh media every 2 days.

### Cell Viability detection(CCK8)

To determine the optimal concentration of hemin to mimic ICH in vitro, we detected the cell viability at 24 hours after hemin exposure. Different concentration of hemin solution and rtPA solution were prepared by dissolving hemin (Sigma, USA) and rtPA (Boehringer Ingelheim) in PBS solution. After discarding the medium, cells were treated with 100 μL specific concentrations (0, 10, 20, 30, 40, 50, 60, 80, 100, 200 μM) of hemin solution and specific concentration (0, 20, 40, 60, 80, 100, 200, 300, 400, 500 nM) of rtPA solution. After 24 h of incubation, the hemin-containing and rtPA-containing medium were discarded, 10 μL CCK-8 solution (Dojindo, Japan) was added to each well and incubated with the cells at 37°C for 30 minutes. Measurement of the absorbance of each well was performed at 450 nm with a microplate reader.

### RNA isolation and library preparation

RNAseq libraries were established from total RNA isolated from primary cortex neuron under different treatment. RNA integrity was assessed using an Agilent 2100 bioanalyser (Agilent Technologies, USA). Transcriptome sequencing and analysis were performed by Novogene, Ltd. (Beijing, China).

### RNA sequencing and analysis of differentially expressed genes

The libraries were sequenced on an Illumina Novaseq platform, and 150 bp paired-end reads were generated. Reads containing poly-N and low-quality reads were removed to obtain cleaned reads, which were retained for subsequent analyses. The clean reads were mapped to the mm9 mouse reference genome using HISAT2. Fragments per kilobase of transcript per million mapped reads (FPKM) for each gene were generated using Cufflinks, and the read counts of each gene were obtained with HTSeqcount. Differential expression Genes (DEGs) was screened using the R package DESeq2. P-value < 0.05 and fold change > 2 or fold change < 0.5 were set as the thresholds for significant differential expression. ClusterProfiler software was used to the analysis of GO or KEGG enrichment of DEGs.

### Transmission Electron Microscopy

After incubated with different treatment for 24h, the primary cortical neuron cells were washed, trypsinized and resuspended. Then we fixed the cells with electron microscope fixative (Servicebio, China) at room tempurature for 2 hours and transferred the mixture at 4°C overnight. The cells were fixed in 1% osmium tetroxide-PBS solution for 1 h at room temperature. The samples were embedded in Epon812(SPI, USA) after dehydrated through a graded series of ethanol. Ultrathin sections were stained with 2.6% uranyl acetate and lead citrate. The TEM was performed using a HITACHI Transmission Electron (HITACHI, Japan) to collect the corresponding images of autophagosomes and ER.

### ER staining with ER-tracker

ER staining was performed according to the instructions of ER-Tracker™ Red kit (Invitrogen E34250). After treatment, the primary cortical neurons were washed twice with HBSS and then incubated in pre-warmed ER-tracker dye solution (1 μM) for approximately 30 min at 37°C. The cells were then fixed by 4% paraformaldehyde and observed using a confocal microscope (Olympus IX71).

### Immunofluorescence

Primary cortical neuron cells cultured on coverslips were treatment with hemin, rtPA or control for 24 hours. After washing twice in PBS, cells were fixed with 4% paraformaldehyde in PBS, permeabilized with 0.3% Triton-X-100, and reacted with 10% BSA at room temperature for 1 hour. Then the cells were incubated with primary antibody overnight at 4°C. After that, the cells were washed with PBS and incubated with secondary antibody for 1 hour at room temperature. Then added 10 ul DAPI staining solution into the coverslips and incubated for 5 minutes at room temperature. After washing with PBS three times, a drop of anti-fluorescent quench sealer was placed in the center of the coverlips. Immunofluorescence images were acquired by a fluorescence microscope (Olympus BX53). The primary antibodies used include the following: Phospho-PERK (CST, 3179S) Cy3-labeled Goat Anti-Rabbit IgG(H+L) (Servicebio, China) was used as secondary antibodies.

### Statistical Analysis

We used the statistical graphing software GraphPad Prism 9.0 for statistical analysis of the data. Behavioral data were processed and analyzed using two-way analysis of variance and Bonferroni test. Western blot and TUNEL staining data were processed using ImageJ software and statistically analyzed using GraphPad Prism 9.0, and one-way analysis of variance was used to process the data and the Turkey test was performed. Data are represented as mean ± standard error (Mean±SEM), and P < 0.05 represented statistical significance. Confocal laminar scan images of ER-tracker were reconstructed to three dimensions using the Surface function of Imaris software. Relevant images of transcriptomic data analysis were drawn using R language. Heat maps were drawn using R language software for mRNA expression of specific genes (autophagy animal (KEGG: mmu04140), positive regulation of neuron apoptotic process (GO:0043525), positive regulation of response to endoplasmic reticulum stress (GO:1905898) between different sample groups. The data were statistically analyzed and plotted using the statistical graphing software GraphPad Prism 9.0.

## Abbreviations

rtPA: recombinant tissue plasminogen activator
MIS: minimally invasive surgery
AKT: AKT kinases group, protein kinase B
EGF: Epidermal Growth Factor
LRP: lipoprotein receptor-related protein
mTOR: mechanistic target of rapamycin kinase
PI3K: Phosphoinositide 3-kinase
RAP: Receptor-associated protein
TXA: tranexamic acid

## Author Contributions

JJ, PZ and ZT conceived and designed research. JJ and SC conducted experiments. JJ, SC, XW, JY, XL, JW and YL analyzed data. SC and XW wrote the manuscript. All authors read and approved the manuscript.

## Data Availability Statement

The original contributions presented in the study are included in the article, further inquiries can be directed to the corresponding authors.

